# Chitinase 3-like-1 Contributes to Acetaminophen-induced Liver Injury by Promoting Hepatic Platelet Recruitment

**DOI:** 10.1101/2021.04.08.439034

**Authors:** Zhao Shan, Leike Li, Constance Lynn Atkins, Meng Wang, Yankai Wen, Jongmin Jeong, Nicolas F. Moreno, Dechun Feng, Xun Gui, Ningyan Zhang, Chun Geun Lee, Jack Angel Elias, William Lee, Bin Gao, Fong Wilson Lam, Zhiqiang An, Cynthia Ju

## Abstract

Hepatic platelet accumulation contributes to acetaminophen (APAP)-induced liver injury (AILI). However, little is known about the molecular pathways involved in platelet recruitment to the liver and whether targeting such pathways could attenuate AILI. The present study unveiled a critical role of chitinase 3-like-1 (Chi3l1) in hepatic platelet recruitment during AILI. Increased Chi3l1 and platelets in the liver were observed in patients and mice overdosed with APAP. Compared to wild-type (WT) mice, Chi3l1^-/-^ mice developed attenuated AILI with markedly reduced hepatic platelet accumulation. Mechanistic studies revealed that Chi3l1 signaled through CD44 on macrophages to induce podoplanin expression, which mediated platelet recruitment through C-type lectin-like receptor 2. Moreover, APAP treatment of CD44^-/-^ mice resulted in much lower numbers of hepatic platelets and liver injury than WT mice, a phenotype similar to that in Chi3l1^-/-^ mice. Recombinant Chi3l1 could restore hepatic platelet accumulation and AILI in Chi3l1^-/-^ mice, but not in CD44^-/-^ mice. Importantly, we generated anti-Chi3l1 monoclonal antibodies and demonstrated that they could effectively inhibit hepatic platelet accumulation and AILI. Overall, we uncovered the Chi3l1/CD44 axis as a critical pathway mediating APAP-induced hepatic platelet recruitment and tissue injury. We demonstrated the feasibility and potential of targeting Chi3l1 to treat AILI.

## Introduction

Acute liver failure (ALF) is a life-threatening condition of massive hepatocyte injury and severe liver dysfunction that can result in multi-organ failure and death.[1] Acetaminophen (APAP) overdose is the leading cause of ALF in Europe and North America and responsible for more cases of ALF than all other aetiologies combined.[1, 2] It is estimated that each week, more than 50 million Americans use products containing APAP and approximately 30,000 patients are admitted to intensive care units every year due to APAP-induced liver injury (AILI).[1, 3] Although N-acetylcysteine (NAC) can prevent liver injury if given in time, there are still 30% of patients who do not respond to NAC.[4] Thus, identification of novel therapeutic targets and strategies is imperative.

APAP is metabolized predominantly by Cytochrome P450 2E1 (CYP2E1) to a reactive toxic metabolite, N-acetyl-p-benzoquinone imine (NAPQI). NAPQI causes mitochondrial dysfunction, lipid peroxidation and eventually cell death.[5] The initial direct toxicity of APAP triggers the cascades of coagulation and inflammation, contributing to the progression and exacerbation of AILI.[5] In patients with APAP overdose, the clinical observations of thrombocytopenia, reduced plasma fibrinogen levels, elevated thrombin-antithrombin, and increased levels of pro-coagulation microparticles strongly suggest concurrent coagulopathy.[6, 7] Similarly, APAP challenge in mice causes a rapid activation of the coagulation cascade and significant deposition of fibrin(ogen) in the liver.[8–10] With regard to the role of platelets in AILI, it is reported that in mice APAP-induced thrombocytopenia correlates with the accumulation of platelets in the liver and that platelet-depletion significantly attenuates AILI.[11] Two recent studies also demonstrate that persistent platelet accumulation in the liver delays tissue repair after AILI in mice.[10, 12] These findings strongly indicate that hepatic platelet accumulation is a key mechanism contributing to AILI. However, little is known about the underlying molecular mechanism of APAP-induced hepatic platelet accumulation and whether targeting this process could attenuate AILI.

Chi3l1 (YKL-40 in humans) is a chitinase-like soluble protein without chitinase activities.[13] It is produced by multiple cell types, including macrophages, neutrophils, fibroblasts, synovial cells, endothelial cells, and tumor cells.[14, 15] Chi3l1 has been implicated in multiple biological processes including apoptosis, inflammation, oxidative stress, infection, and tumor metastasis.[16] Elevated serum levels of Chi3l1 have been observed in various liver diseases, such as hepatic fibrosis, non-alcoholic fatty liver, alcoholic liver disease, and hepatocellular carcinoma. [13, 17–19] However, the biological function of Chi3l1 in liver disease is not clear. Our previous study revealed an important role of Chi3l1 in promoting intrahepatic coagulation in concanavalin A-induced hepatitis.[20] Given the importance of intrahepatic coagulation in the mechanism of AILI, we wondered whether Chi3l1 is involved in platelets accumulation during AILI.

In the current study, we observed elevated levels of Chi3l1 in patients with APAP-induced acute liver failure and in mice challenged with APAP overdose. Our data demonstrated a central role of Chi3l1 in APAP-induced hepatic platelet recruitment through CD44. Importantly, we found that targeting Chi3l1 by monoclonal antibodies could effectively inhibit platelet accumulation in the liver and markedly attenuate AILI.

## Results

### Chi3l1 is upregulated and plays a critical role in AILI

Although elevated serum levels of Chi3l1 has been observed in chronic liver diseases,[13, 17–19] modulations of Chi3l1 levels during acute liver injury have not been reported. Our data demonstrated, for the first time, that compared with healthy individuals, patients with AILI displayed higher levels of Chi3l1 in the liver and serum (Figures 1A, B). Similarly, in mice treated with APAP, hepatic mRNA and serum protein levels of Chi3l1 were upregulated (Figures 1C, D). To determine the role of Chi3l1 in AILI, we treated wild-type (WT) mice and Chi3l1-knockout (Chi3l1^-/-^) mice with APAP. Compared with WT mice, serum ALT levels and the extent of liver necrosis were dramatically lower in Chi3l1^-/-^ mice (Figures 1E, F). Moreover, administration of recombinant mouse Chi3l1 protein (rmChi3l1) to Chi3l1^-/-^ mice enhanced liver injury to a similar degree observed in APAP-treated WT mice (Figures 1E, F). These data strongly suggest that Chi3l1 contributes to AILI.

**Figure 1.**
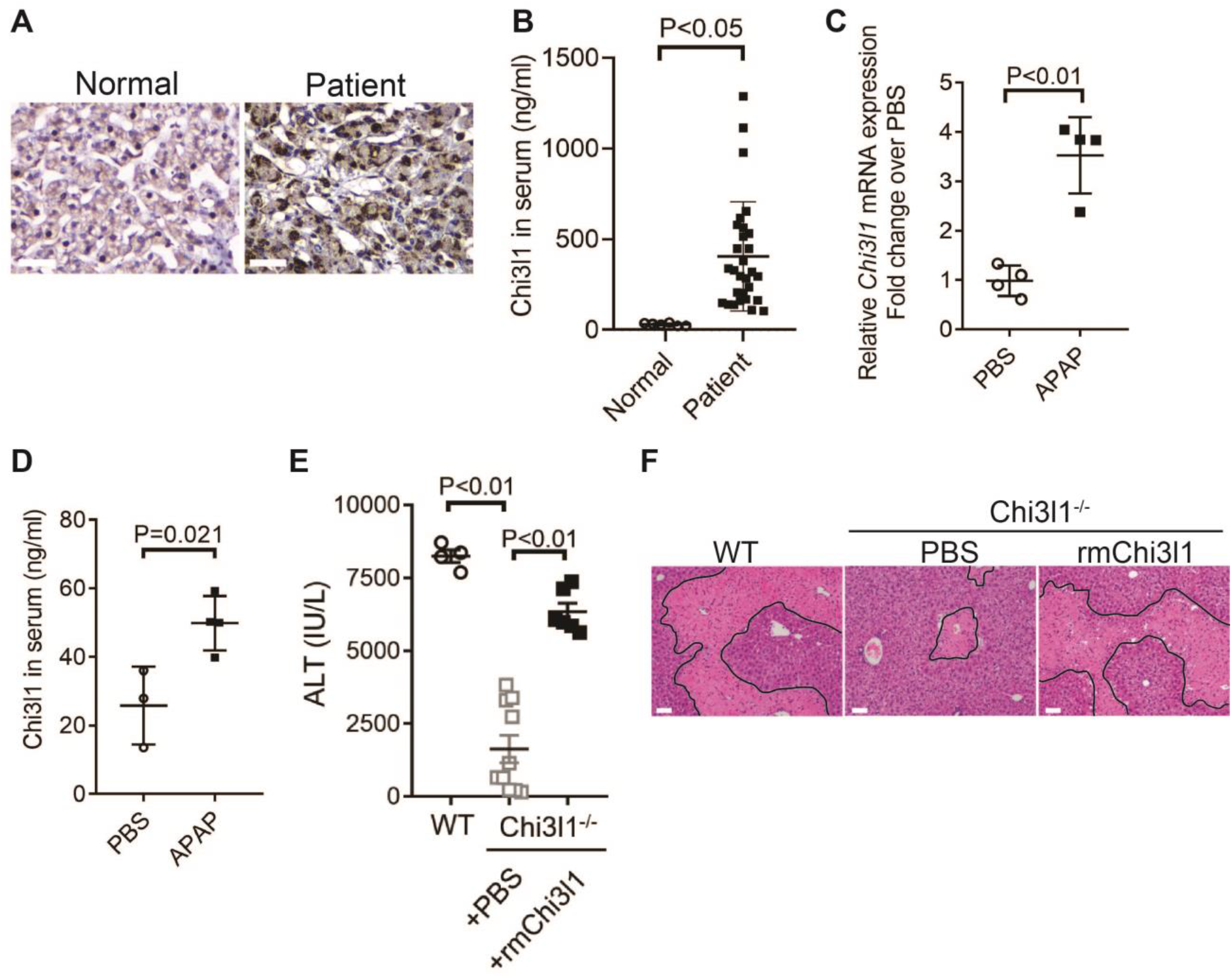
Chi3l1 is upregulated and play a critical role in AILI. (**A**) IHC staining for Chi3l1 in normal liver biopsies (Normal) and those from patients with AILI (Patient). Images shown are representative of 10 samples/group. Scale bar, 250μm. (**B**) ELISA analysis of Chi3l1 in serum of healthy individuals (Normal, n=6) and those from patients with AILI (Patient, n=29). Data were presented as median + interquartile range. (**C, D**) Male C57B/6 mice treated with PBS or APAP. (**C**) *Chi3l1* mRNA in liver homogenates and (**D**) Chi3l1 protein levels in serum were measured by qRT-PCR and ELISA at 3hrs and 24hrs, respectively (n=4 mice/group). (**E, F**) Male C57B/6 (WT) and Chi3l1^-/-^ mice were treated with APAP. Additionally, Chi3l1^-/-^ mice were divided into two groups treated with either PBS or recombinant mouse Chi3l1 (rmChi3l1) simultaneously with APAP (n=4-10 mice/group). (**E**) Serum levels of ALT and (**F**) liver histology with necrotic areas outlined were evaluated 24hrs after APAP treatment. Scale bar, 250μm. Mann-Whitney test was performed in **B**. Two-tailed, unpaired student t-test was performed in **C, D**. One-way ANOVA were performed in **E**.

### Chi3l1 contributes to AILI by promoting hepatic platelet recruitment

Thrombocytopenia is often observed in patients with APAP overdose.[6, 7, 21] We hypothesized that this phenomenon may be attributed to the recruitment of platelets into the liver. We performed immunohistochemical (IHC) staining of liver biopsies from patients with APAP-induced liver failure and found markedly increased numbers of platelets compared with normal liver tissues (Figure 2A). Similarly, in mice treated with APAP, a marked increase of platelets in the liver was observed by intravital microscopy (Figure 2B). It is reported that depletion of platelets prior to APAP treatment can prevent liver injury in mice.[11] Our data demonstrated that even after APAP treatment, depletion of platelets could still attenuate AILI (Figures 2C, D; Supplementary Figure 1). These data indicate a critical contribution of platelets to AILI. Given the role of Chi3l1 in promoting intrahepatic coagulation in concanavalin A-induced hepatitis,[20] we hypothesized that Chi3l1 might be involved in platelet recruitment to the liver during AILI. To examine this hypothesis, we detected platelets in the liver by IHC using anti-CD41 antibody. Comparing with WT mice, we observed much fewer platelets in the liver after APAP treatment (Figure 2E). Moreover, administration of rmChi3l1 to Chi3l1^-/-^ mice restored hepatic platelet accumulation similar to APAP-treated WT mice (Figure 2E). These data suggest that Chi3l1 plays a critical role in promoting hepatic platelet accumulation, thereby contributing to AILI.

**Figure 2.**
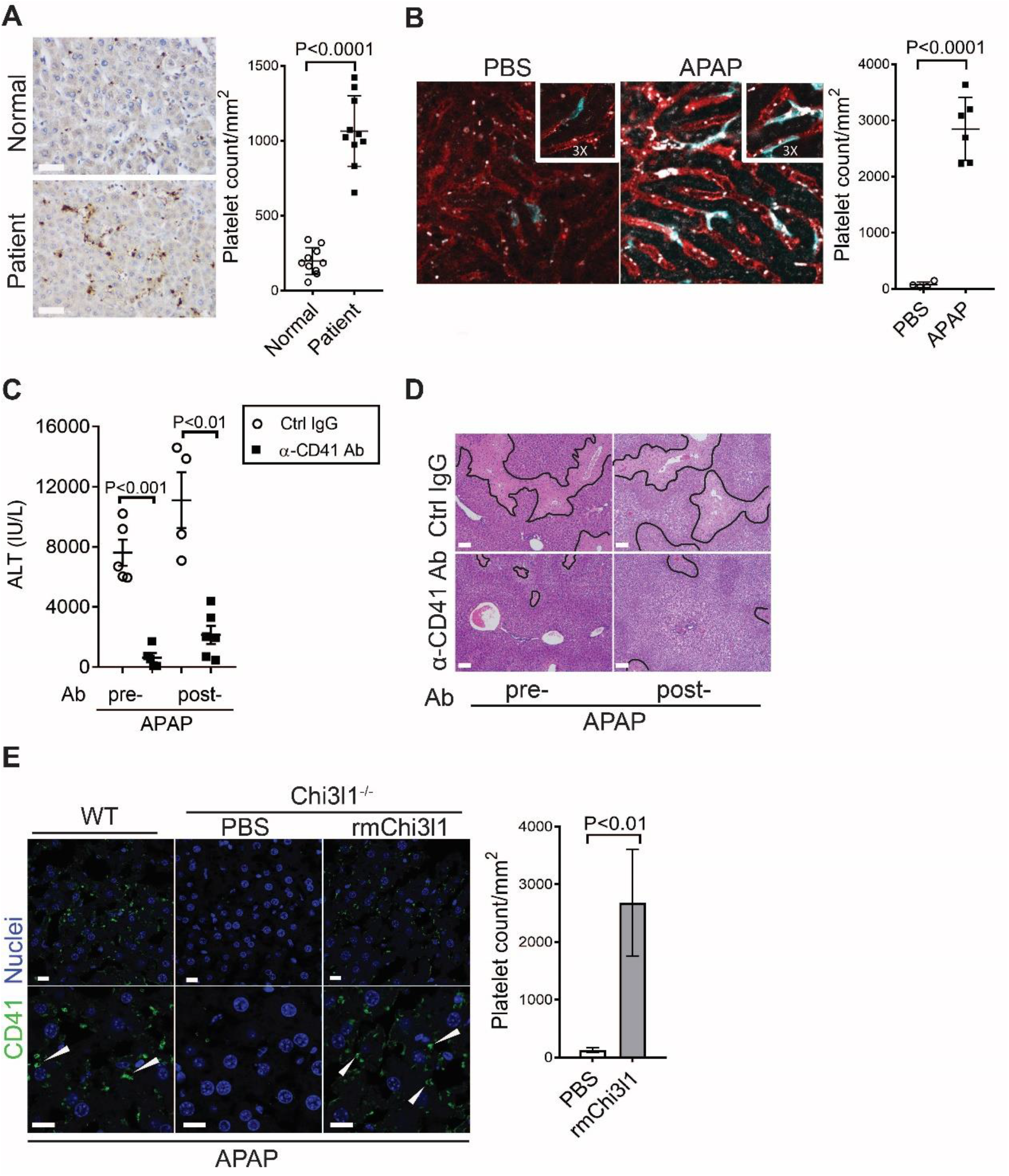
Chi3l1 contributes to AILI by promoting hepatic platelet recruitment. (**A**) IHC staining to detect platelets (CD41^+^) in healthy liver biopsies (Normal) and those from patients with AILI (Patient). Scale bar, 250μm. (n=10/group) (**B**) Male C57B/6 mice treated with PBS or APAP. Intravital microscopy analyses were performed around 3 hrs post APAP. Mϕs (cyan) and platelets (white) in liver sinusoids (red) are indicated. Representative images were chosen from intravital microscopy videos: https://bcm.box.com/s/15hmtryyrdl302mihrsm034ure87x4ea (Supplemental video 1, PBS treatment) and https://bcm.box.com/s/tuljfmstvv4lvoksx16fkxkpirkekynz (Supplemental Video 2, APAP treatment)(n=6-7 mice/group, 4-15 videos/mouse). Two-tailed, unpaired student t-test was performed. (**C-E**) Male C57B/6 (WT) mice were treated with control IgG (Ctrl IgG) or an anti-CD41 antibody (α-CD41 Ab) either 3hrs before or 3hrs after APAP administration. (**C**) Serum levels of ALT and (**D**) liver histology with necrotic areas outlined were evaluated 24hrs after APAP treatment (n=5 mice/group in C, D). Scale bar, 250μm. (**E**) IF staining was performed to detect intrahepatic platelets (CD41^+^) 3hrs after APAP treatment (n=3 mice/group). Scale bar, 25μm. Two-tailed, unpaired student t-test was performed in **A-C, E.**

### Chi3l1 functions through its receptor CD44

To further understand how Chi3l1 is involved in platelet recruitment, we set out to identify its receptor. We isolated non-parenchymal cells (NPCs) from WT mice at 3h after APAP treatment and incubated the cells with His-tagged rmChi3l1. The cell lysate was subjected to immunoprecipitation using an anti-His antibody. The “pulled down” fraction was subjected to LC/MS analyses, and a partial list of proteins identified is shown in Supplementary Table 1. Among the potential binding proteins, we decide to further investigate CD44, which is a cell surface receptor expressed on diverse mammalian cell types, including endothelial cells, epithelial cells, fibroblasts, keratinocytes and leukocytes.[22] Immunoprecipitation experiments using liver homogenates from APAP-treated WT and CD44^-/-^ mice demonstrated that the anti-CD44 antibody could “pull down” Chi3l1 from WT but not CD44^-/-^ liver homogenates (Figure 3A). Supporting this finding, interferometry measurements using recombinant human Chi3l1 (rhChi3l1) revealed a direct interaction between Chi3l1 and CD44 (Kd = 251nM, Figure 3B). Moreover, we incubated rhChi3l1 with human CD44 and then performed immunoprecipitation with an anti-CD44 antibody. Data shown in Figure 3C confirmed that Chi3l1 directly binds to CD44. Together, these results suggest that CD44 is a receptor for Chi3l1.

**Figure 3.**
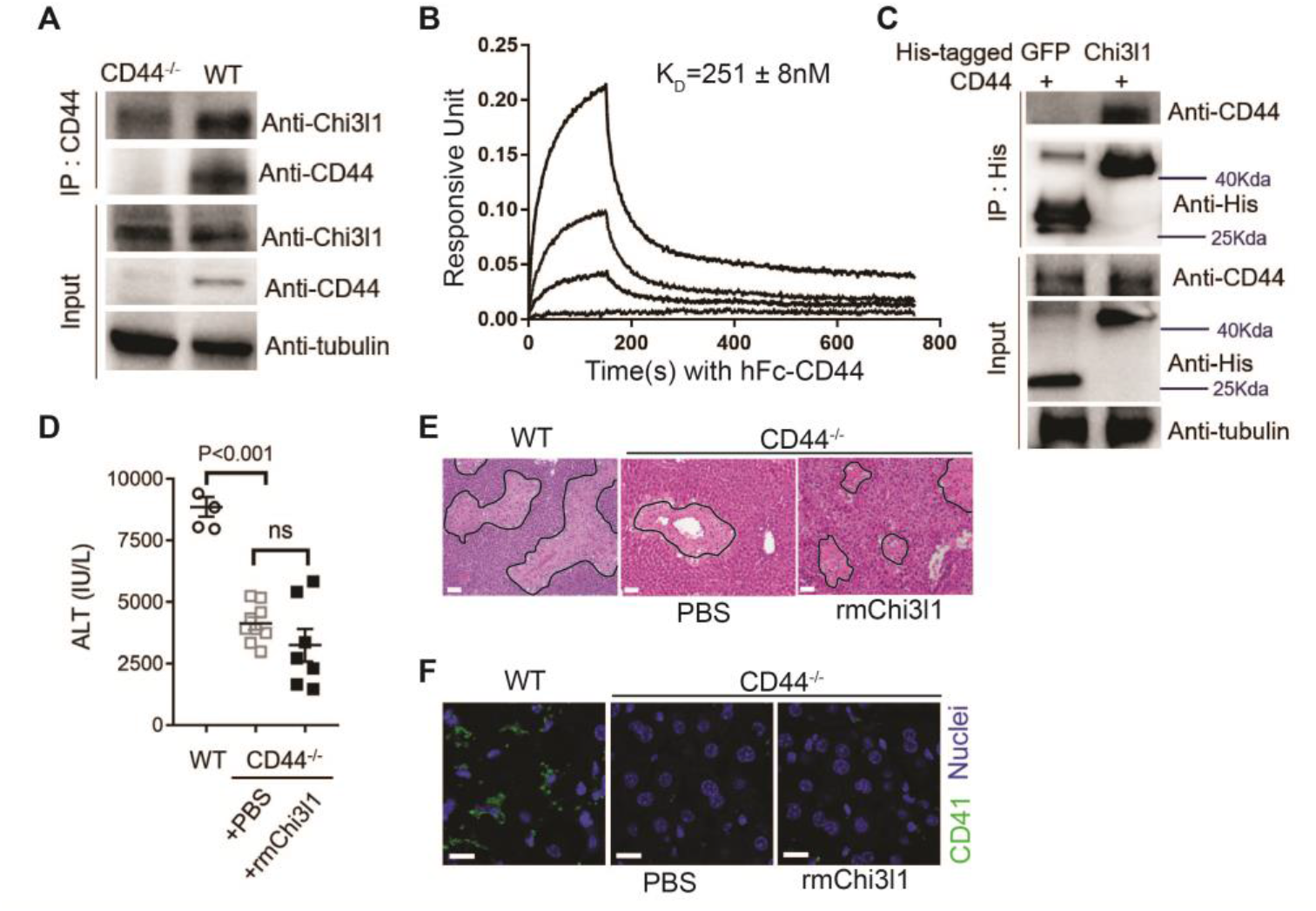
Chi3l1 functions through its receptor CD44. (**A**) Immuno-precipitation with anti-CD44 antibody was performed using liver homogenates obtained from WT and CD44^-/-^ mice treated with APAP for 2hrs. Input proteins and immune-precipitated proteins were blotted with the indicated antibodies. (**B**) Interferometry measurement of the binding kinetics of human His-Chi3l1 with human Fc-CD44. (**C**) His-tagged control GFP and human Chi3l1 were incubated with recombinant human CD44. Proteins bound to Chi3l1 were immune-precipitated with an anti-His antibody. Input proteins and immune-precipitated proteins were blotted with indicated antibodies. (**D-F**) Male WT mice were treated with APAP and CD44^-/-^ mice were treated with PBS or rmChi3l1 plus APAP. (**D**) Serum levels of ALT and (**E**) liver histology with necrotic areas outlined were evaluated 24hrs after APAP treatment (n=4-9 mice/group in A, B). Scale bar, 250μm. (**F**) IF staining was performed to detect intrahepatic platelets (CD41^+^) 3hrs after APAP treatment (n=3 mice/group). Scale bar, 25μm. One-way ANOVA were performed in **D.**

To investigate the role of CD44 in mediating the function of Chi3l1, we treated CD44^-/-^ mice with rmChi3l1 simultaneously with APAP challenge. We found that rmChi3l1 had no effect on platelet recruitment or AILI in CD44^-/-^ mice (Figures 3D-F). This is in stark contrast to restoring platelet accumulation and increasing AILI by rmChi3l1 treatment in Chi3l1^-/-^ mice (Figure 1E, F; 2E). However, these effects of rmChi3l1 in Chi3l1^-/-^ mice were abrogated when CD44 was blocked by using an anti-CD44 antibody (Supplementary Figures 2A-C). Together, these data demonstrate a critical role of CD44 in mediating Chi3l1-induced hepatic platelet accumulation and AILI.

CYP2E1-mediated APAP bio-activation to form N-acetyl para quinoneimine (NAPQI) and the detoxification of NAPQI by glutathione (GSH) are important in determining the degrees of AILI.[5] Although unlikely, there is a possibility that the phenotypes observed in Chi3l1^-/-^ and CD44^-/-^ mice were due to the effects of gene deletion on APAP bio-activation. To address this concern, we compared the levels of GSH, liver CYP2E1 protein expression, and NAPQI-protein adducts among WT, Chi3l1^-/-^ and CD44^-/-^ mice (Supplementary Figures 3A-C). However, we did not observe any difference, suggesting that Chi3l1 or CD44 deletion does not affect APAP bio-activation and its direct toxicity to hepatocytes.

### Hepatic Mϕs promote platelet recruitment

To further identify the cell type on which Chi3l1 binds to CD44, we incubated liver NPCs with His-tagged rmChi3l1. We found that almost all CD44^+^Chi3l1^+^ cells were F4/80^+^ Mϕs (Supplemental Figure 2D). This finding suggested the possible involvement of hepatic Mϕs in platelet recruitment. We performed IHC staining of liver biopsies from AILI patients and observed co-localization of Mϕs (CD68+) and platelets (CD41+) (Figure 4A). In the livers of APAP-treated mice, adherence of platelets to Mϕs was also observed by IHC (Figure 4B) and intravital microscopy (Figure 2B). Quantification of the staining confirmed that there were higher numbers of platelets adherent to Mϕs than to LSECs after APAP challenge (Figure 4B).

**Figure 4.**
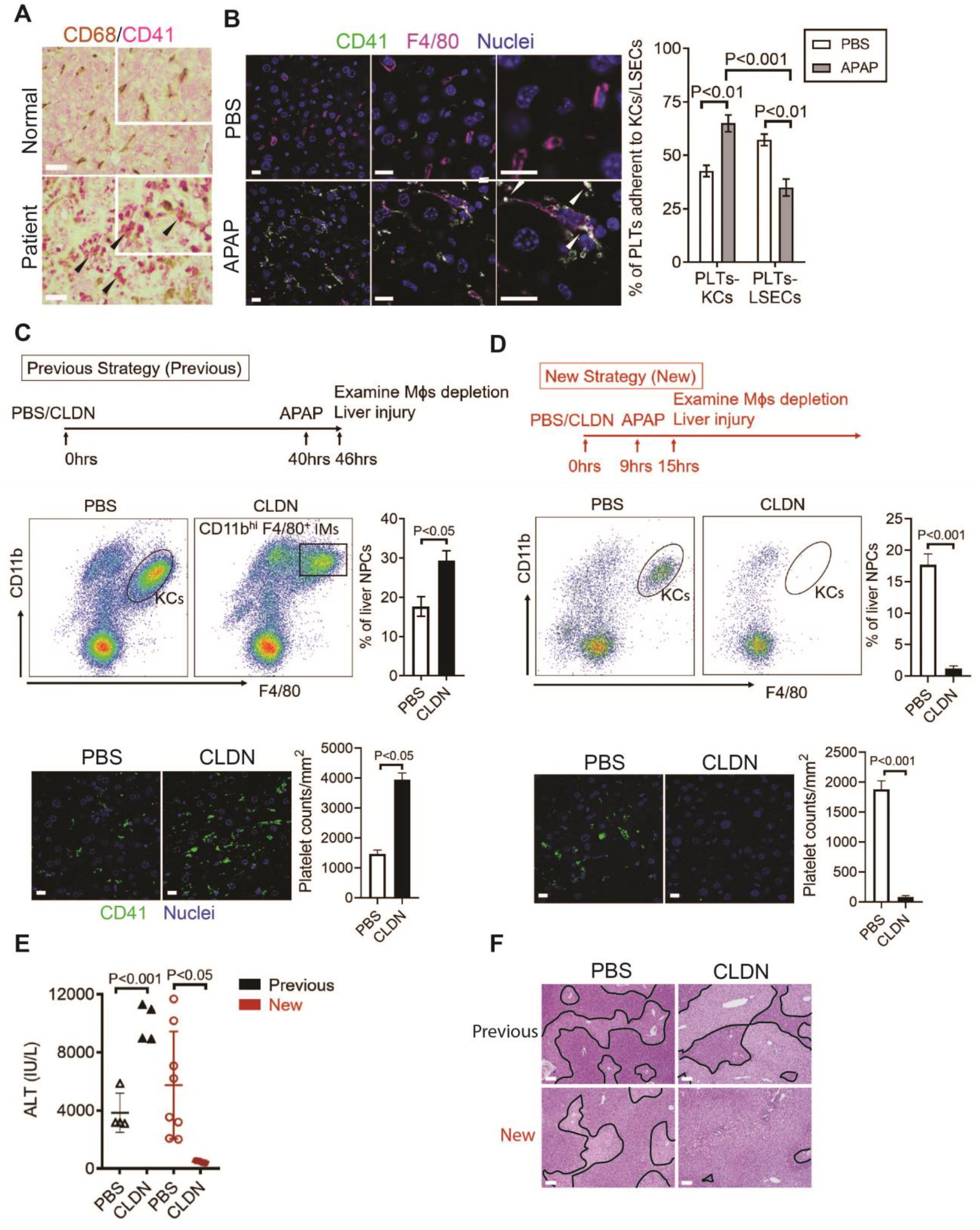
Hepatic Mϕs promote platelet recruitment. (**A**) IHC staining for macrophages (CD68^+^) and platelets (CD41^+^) in normal liver biopsies (Normal) and those from patients with AILI (Patient) (n=10/group). Scale bar, 25μm. (**B**) IF staining for intrahepatic platelets (CD41^+^) and KCs (F4/80^+^) in male C57B/6 mice treated with PBS or APAP for 3hrs. Scale bar, 25μm. Arrowheads indicate platelets adherent to KCs. Quantification of platelets adherent to KCs or LSECs. (**C-F**) Male C57B/6 mice were injected with either empty liposomes containing PBS (PBS) or liposomes containing clodronate (CLDN), followed by APAP treatment. (**C, D**) NPCs were isolated and underwent flow cytometry analysis. Indicated cells were gated on single live CD45^+^CD146^-^ cells. IF staining was performed to detect intrahepatic platelets (CD41^+^). Scale bar, 25μm. (**E**) Serum levels of ALT and (**F**) liver histology with necrotic areas outlined. Scale bar, 250μm. (n=6 mice/group in B-F). Two-tailed, unpaired student t-test was performed in **B-D, F**.

To further investigate the role of hepatic Mϕs in platelet recruitment during AILI, we performed Mϕ-depletion experiments using liposome-encapsulated clodronate (CLDN). We first followed a previously published protocol[23–25]and injected CLDN around 40hrs prior to APAP treatment (Figure 4C, “Previous Strategy”). We examined the efficiency of Mϕ-depletion by flow cytometry analysis, which can distinguish resident Kupffer cells (KCs, CD11b^low^F4/80^+^) from infiltrating Mϕs (IMs, CD11b^hi^F4/80^+^).[26] We found that compared with control mice treated with empty liposomes, there were actually more Mϕs, consisted of mainly IMs, in the liver of CLDN-treated mice (Figure 4C). Consistent with the increase of Mϕs, there were also higher numbers of platelets in the liver of CLDN-treated mice (Figure 4C). These findings suggest that although KCs are depleted using the “Previous Strategy”, the treatment of CLDN induces the recruitment of IMs, resulting in higher numbers of Mϕs in the liver at the time of APAP treatment. As reported, this treatment strategy resulted in exacerbated AILI (Fig. 4E, F “Previous Strategy”), which had led to the conclusion in published reports that KCs play a protective role against AILI.[23–25] However, alternatively the enhanced injury could be due to increased IMs and platelet accumulation.

To better investigate the role of hepatic Mϕs in platelet recruitment, we set out to identify a time period in which both KCs and IMs are absent after CLDN treatment. We measured hepatic Mϕs by flow cytometry at varies times after CLDN treatment and established a “New Strategy”, in which mice were injected with CLDN and after 9hrs treated with APAP. As shown in Fig. 4D, at 6hrs after APAP challenge (15hrs after CLDN), both KCs and IMs were dramatically reduced. Interestingly, when compared to control mice treated with empty liposomes, CLDN-treated mice developed markedly reduced liver injury with nearly no platelet accumulation in the liver (Figures 4D-F “New Strategy”). These data suggest that hepatic Mϕs play a crucial role in platelet recruitment into the liver, thereby contributing to AILI.

### Chi3l1/CD44 signaling in Mϕs upregulates podoplanin expression and platelet adhesion

To further understand how Chi3l1/CD44 signaling in Mϕs promotes platelet recruitment, we measured Mϕs expression of a panel of adhesion molecules known to be important in platelet recruitment.[27–30] Our data showed that podoplanin is expressed at a much higher level in hepatic Mϕs isolated from APAP-treated WT mice than those from Chi3l1^-/-^ or CD44^-/-^ mice (Figure 5A). Interestingly, rmChi3l1 treatment of Chi3l1^-/-^, but not CD44^-/-^ mice, markedly increased the podoplanin mRNA and protein expression levels in Mϕs (Figures 5B, C). To examine the role of podoplanin in mediating platelet adhesion to Mϕs, we blocked podoplanin using an anti-podoplanin antibody in Chi3l1^-/-^ mice reconstituted with rmChi3l1. As shown in Figures 5D-F, blockade of podoplanin not only abrogated rmChi3l1-mediated platelet recruitment into the liver, but also significantly reduced its effect on increasing AILI in Chi3l1^-/-^ mice.

**Figure 5.**
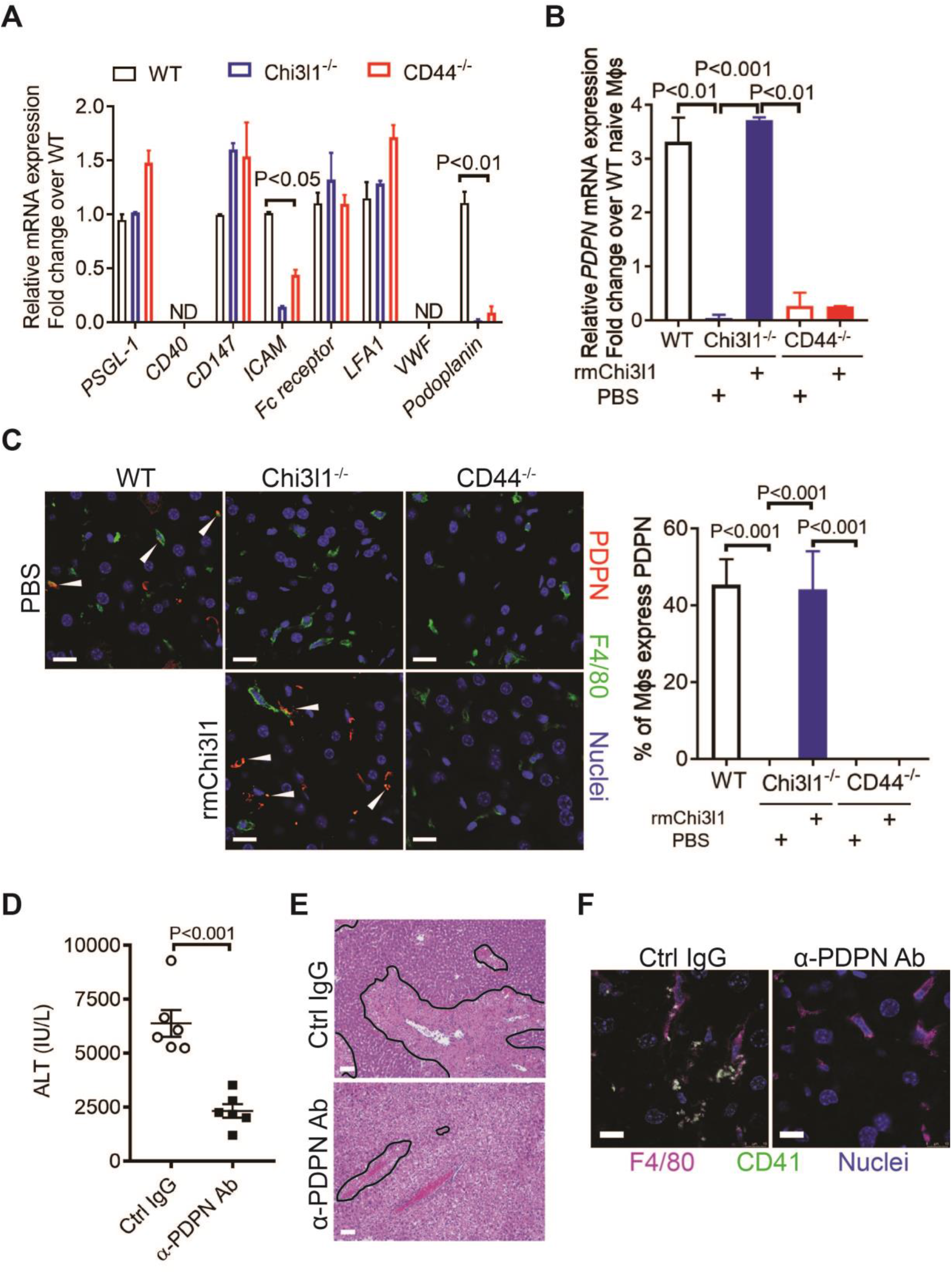
Chi3l1/CD44 signaling in Mϕs upregulates podoplanin expression and platelet adhesion. (**A**) Male WT, Chi3l1^-/-^, CD44^-/-^ mice were treated with APAP (n=4 mice/group). After 3hrs, mice were sacrificed and Mϕs were isolated to measure mRNA levels of various adhesion molecules, including P-Selectin Glycoprotein Ligand 1 (PSGL-1), CD40, CD147, Fc receptor, intercellular adhesion molecule (ICAM), lymphocyte function-associated antigen (LFA1), von Willebrand Factor (VWF), and podoplanin. One-way ANOVA were performed. (**B, C**) WT mice were treated with APAP. Chi3l1^-/-^ and CD44^-/-^ mice were treated with PBS or rmChi3l1 followed by APAP challenge simultaneously and mice were sacrificed 3hrs after APAP (n=3 mice/group). (**B**) Mϕs were isolated and mRNA levels of *PDPN* in Mϕs were analyzed by qRT-PCR. (**C**) IF staining of liver sections for PDPN and F4/80 is shown and the proportions of Mϕs that express PDPN were quantified, Scale bar, 25μm. (**D-F**) Chi3l1^-/-^ mice reconstituted with rmChi3l1 were treated with either Ctrl IgG or α-podoplanin Ab for 16hrs and subsequently challenged with APAP. (**D**) Serum levels of ALT and (**E**) liver histology were evaluated 24hrs after APAP treatment (n=6 mice/group). Scale bar, 250μm. (**F**) IF staining for intrahepatic platelets (CD41^+^) and Mϕs (F4/80+) was performed 3hrs after APAP (n=3 mice/group). Scale bar, 25μm. One-way ANOVA were performed in **A-C**. Two-tailed, unpaired student t-test was performed in **E**.

C-type lectin-like receptor 2 (Clec-2) is the only platelet receptor known to bind podoplanin[31]. To further elucidate the role of podoplanin in mediating platelet adhesion to Mϕs, we isolated Mϕs from WT mice treated with APAP. After treating Mϕs with anti-podoplanin antibody or IgG as control, we added platelets. Immunofluorescence staining of podoplanin and Clec-2 showed that the Clec-2-expressing platelets only bound to IgG-treated, but not anti-podoplanin-treated Mϕs (Supplementary Figure 4). Together, our data demonstrate that Mϕs recruit platelets through podoplanin and Clec-2 interaction, and that the podoplanin expression on Mϕs is regulated by Chi3l1/CD44 signaling.

### Evaluation of the therapeutic potential of targeting Chi3l1 in the treatment of AILI

Although NAC greatly reduces morbidity and mortality from ALF due to APAP overdose, the death rate and need for liver transplantation remain unacceptably high. While elucidating the underlining biology of Chi3l1 in AILI, we also generated monoclonal antibodies specifically recognizing either mouse or human Chi3l1. We screened a panel of anti-mouse Chi3l1 monoclonal antibodies (α-mChi3l1 mAb) to determine their efficacies in attenuating AILI. We injected WT mice with an α-mChi3l1 mAb or IgG at 3h after APAP challenge. Our data showed that clone 59 (C59) had the most potent effects on inhibiting APAP-induced hepatic platelet accumulation and attenuating AILI (Figures 6A-C).

**Figure 6.**
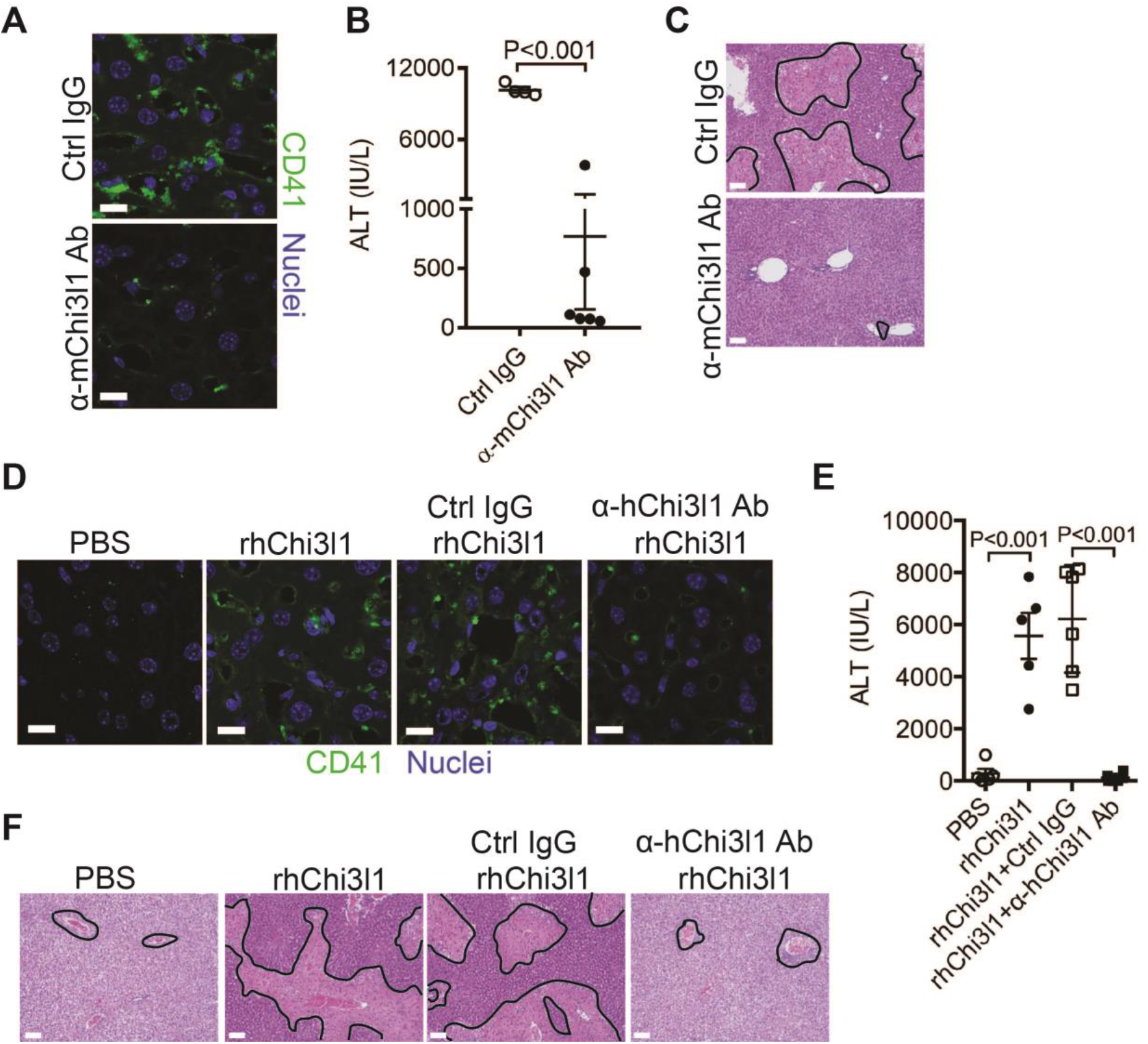
Evaluation of the therapeutic potential of targeting Chi3l1 in the treatment of AILI. (**A-C**) Male C57B/6 mice were treated with APAP for 3hrs, followed by *i.p*. injection of either a control IgG (Ctrl IgG) or an anti-mouse Chi3l1 Ab (α-mChi3l1 Ab, C59). (**A**) IF staining for intrahepatic platelets (CD41^+^) was performed 6hrs after APAP treatment (n=3 mice/group). Scale bar, 25μm. (**B**) Serum levels of ALT and (**C**) liver histology were evaluated 24hrs after APAP treatment (n=4-6 mice/group). Scale bar, 250μm. (**D-F**) Chi3l1^-/-^ mice were treated with APAP plus PBS or recombinant human Chi3l1 (rhChi3l1) for 3hrs as indicated and APAP plus rhChi3l1 treatment group were either without treatment or treated with a control IgG (Ctrl IgG) or an anti-human Chi3l1 Ab (α-hChi3l1 Ab, C7). (**D**) IF staining was performed to identify intrahepatic platelets (CD41^+^) 6hrs after APAP treatment. Scale bar, 25μm. (**E**) Serum levels of ALT and (**F**) liver histology were evaluated 24hrs after APAP treatment. Scale bar, 250μm. (n=5-10 mice/group in **D-F**). Two-tailed, unpaired student t-test was performed in **B**. One-way ANOVA were performed in **F.**

To evaluate the potential of targeting Chi3l1 as a treatment for AILI in humans, we screened all of the α-hChi3l1 mAb we generated by IHC staining of patients’ liver biopsies (data not shown) and selected the best clone for *in vivo* functional studies. Because the amino acid sequence homology between human and mouse Chi3l1 is quite high (76%), we treated Chi3l1^-/-^ mice with rhChi3l1. We found that rhChi3l1 was as effective as rmChi3l1 in promoting platelet recruitment and increasing AILI in Chi3l1^-/-^ mice (Figures 6D-F). To our excitement, the α-hChi3l1 mAb treatment could abrogate platelet recruitment and dramatically reduce liver injury (Figures 6D-F). Together, these data indicate that monoclonal antibody-based blocking of Chi3l1 may be an effective therapeutic strategy to treat AILI, and potentially other acute liver injuries.

## Discussion

The current study unveiled an important function of Chi3l1 in promoting platelet recruitment into the liver after APAP overdose, thereby playing a critical role in exacerbating APAP-induced coagulopathy and liver injury. Our data demonstrate that Chi3l1 signals through CD44 on Mϕs to upregulate podoplanin expression and promote platelet recruitment (Figure 7). Moreover, we report for the first time significant hepatic accumulation of platelets and marked upregulation of Chi3l1 in patients with ALF caused by APAP overdose. Importantly, we demonstrate that neutralizing Chi3l1 with monoclonal antibodies can effectively inhibit hepatic platelet accumulation and mitigate liver injury caused by APAP, supporting the potential and feasibility of targeting Chi3l1 as a therapeutic strategy to treat AILI.

**Figure 7.**
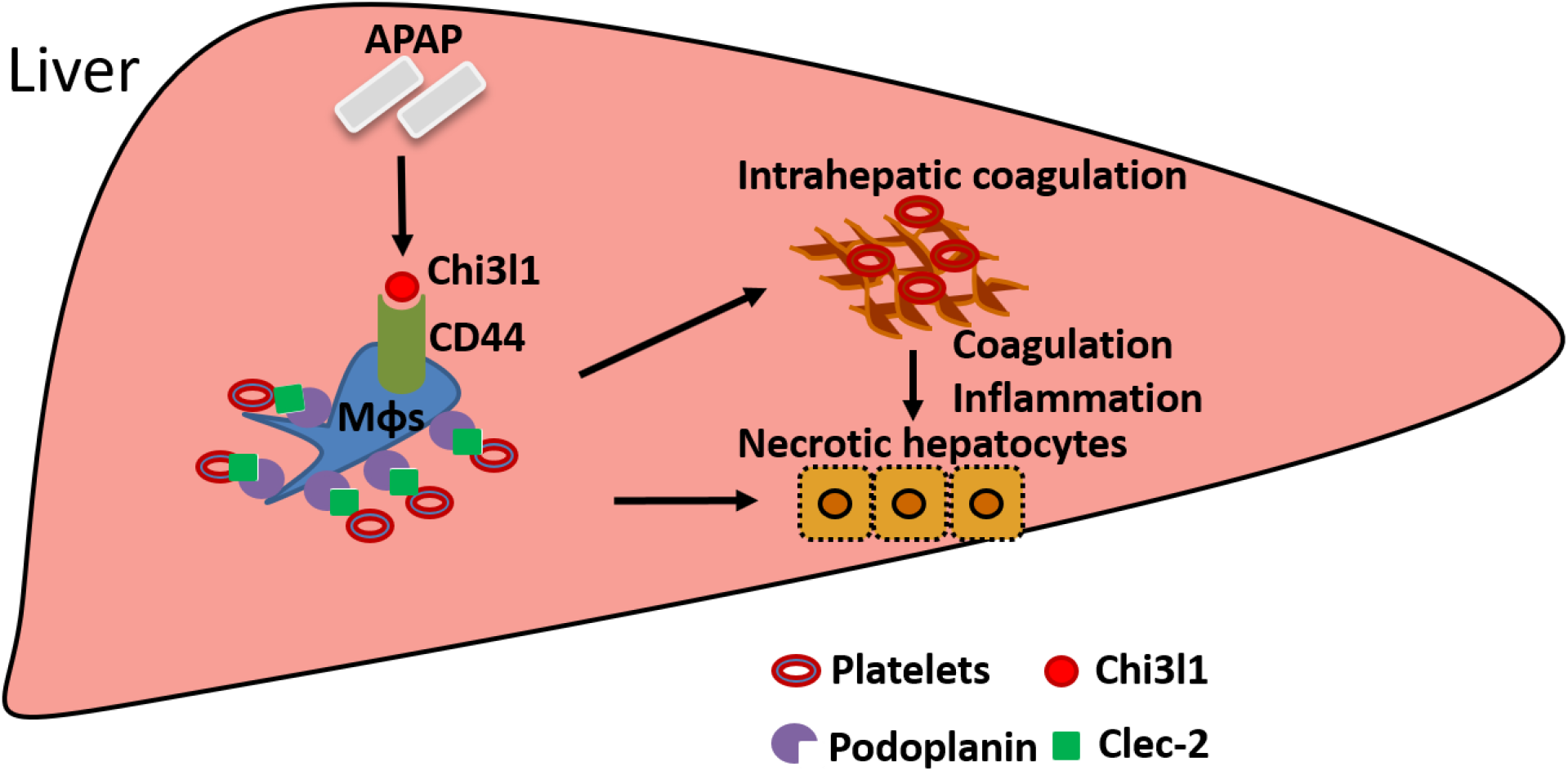
Schematic summary of the main findings. APAP overdose induces Chi3l1 expression, which binds CD44 on Mϕs and promotes Mϕs-mediated platelets recruitment through podoplanin/Clec-2 interaction. Recruited platelets further contribute to AILI.

The elevation of serum levels of Chi3l1 has been observed in various liver diseases, [13, 17–19] but studies of its involvement in liver diseases have only begun to emerge. There are several reports describing a role of Chi3l1 in models of chronic liver injuries caused by alcohol, CCl4 or high-fat diet. [32–35] However, the molecular and cellular mechanisms accounting for the involvement of Chi3l1 have yet to be defined. The present study unveils a function of Chi3l1 in promoting platelet recruitment to the liver during acute injury. We provide compelling data demonstrating that Chi3l1, acting through its receptor CD44 on Mϕs to recruit platelets, thereby contributing to AILI. Multiple receptors of Chi3l1 have been identified, including IL-13Rα2, CRTH2, TMEM219, and galectin-3. [36–40] The fact that Chi3l1 could bind to multiple receptors is consistent with a diverse involvement of Chi3l1 under different disease contexts. A recent study showed that Chi3l1 was upregulated during gastric cancer (GC) development and that through binding to CD44, it activated Erk, Akt, and β-catenin signaling, thereby enhancing GC metastasis. [39] Our studies illustrated a novel role of Chi3l1/CD44 interaction in the recruitment of hepatic platelets and contribution to AILI. Our in vivo studies using CD44^-/-^ mice and anti-CD44 antibody provide strong evidence that CD44 mediates the effects of Chi3l1. Our observation that Chi3l1 predominantly binds to CD44 on Mϕs, but not other CD44-expressing cells in the liver, suggests two possibilities which warrant further investigation. First, Chi3l1 may bind a specific isoform of CD44 that is uniquely expressed by Mϕs. Second, the Chi3l1-CD44 interaction requires binding of a co-receptor, which is expressed on Mϕs but not on other CD44-expressing cells in the liver.

We identified hepatic Mϕs as a key player in promoting platelet recruitment to the liver during AILI. Given the involvement of platelets in AILI, this finding would suggest that hepatic Mϕs also contribute to liver injury. The role of hepatic Mϕs in AILI has been a topic of debate and the current understanding is confined by the limitation of the methods used to deplete these cells. [23-25, 41, 42] Several previous studies using CLDN to deplete Mϕs concluded that these cells play a protective role against AILI. [23–25] However, in those studies, Mϕ-depletion was confirmed by IHC staining of F4/80, which cannot distinguish KCs from IMs. Our laboratory and others had since developed a flow cytometric approach to detect and distinguish the two Mϕs populations. Using flow cytometry to monitor Mϕ-depletion, we found that the timing of CLDN treatment was critical. In the previously published reports, mice were treated with CLDN around 2 days before APAP challenge. [23–25] Using this treatment regimen, IMs became abundant prior to APAP treatment, even though KCs were depleted. Without this knowledge, previous studies attributed the worsened AILI to the depletion of KCs. However, the advancement of knowledge on the recruitment of IMs and their contribution to acute liver injury offers an alternative interpretation that the worsened AILI is due to IM accumulation.[12, 26, 43, 44] In the current study, we analyzed KCs and IMs in the liver at various time points after CLDN treatment to identify a new strategy to achieve more complete hepatic Mϕ-depletion. Our data demonstrated that when both Mϕs populations were absent at the time of APAP treatment, platelet recruitment was abrogated and AILI was significantly reduced. During the preparation of this manuscript, a study was published describing that IMs could recruit platelets.[12] Together, these data suggest that hepatic Mϕs (both KCs and IMs) play a crucial role in promoting hepatic platelet accumulation, thereby contributing to AILI.

Our data suggest that platelet-derived Clec-2 interacts with podoplanin expressed on Mϕs, resulting in platelet recruitment to the liver during the early phase of AILI. The role of podoplanin/Clec-2 interaction in platelet recruitment and thromboinflammation has been indicated in multiple inflammatory and infectious conditions.[12, 30, 31] Our data, for the first time, provide evidence that the podoplanin expression on Mϕs is regulated by the Chi3l1/CD44 axis. Future studies focusing on gaining molecular insight into such regulation are warranted. An increasing number of studies suggest that platelets play an important, but paradoxical role in liver injury. It has been proposed that they contribute to tissue damage during injury phase but promote tissue repair at later time points.[45] However, two recent studies of AILI demonstrate that persistent platelet accumulation in the liver significantly delays liver repair. One study described a podoplanin/Clec-2 interaction between platelets and hepatic IMs during tissue repair, and demonstrated a detrimental role of such interaction through blocking the recruitment of reparative neutrophils.[12] Another study showed that AILI was associated with elevated plasma levels of von Willebrand Factor (vWF), which prolonged hepatic platelet accumulation and delayed repair of APAP-injured liver in mice.[10] These studies together with our finding that platelets drive tissue damage during early stage of AILI suggest that platelets may be a therapeutic target to treat acute liver injury.

Our studies uncovered a previously unrecognized involvement of the Chi3l1/CD44 axis in AILI and provided insights into the mechanism by which Chi3l1/CD44 signaling promotes hepatic platelet accumulation and liver injury after APAP challenge. Taking our findings one-step further toward clinical application, we demonstrated the feasibility of targeting Chi3l1 by mAbs to attenuate AILI. There is an unmet need for developing treatments for AILI, as NAC is the only antidote at present. However, the efficacy of NAC declines rapidly when initiated more than a few hours after APAP overdose, long before patients are admitted to the clinic with symptoms of severe liver injury.[46] Our studies provide strong support for the potential targeting of Chi3l1 as a novel therapeutic strategy to improve the clinical outcomes of AILI and perhaps other acute liver injury conditions.

## Methods

### Animal experiments and procedures

C57BL/6J and CD44^-/-^ mice were purchased from the Jackson Laboratory. Chi3l1^-/-^ mice were provided by Dr. Jack Elias (Brown University, Providence, RI, United States). All mouse colonies were maintained at the animal core facility of University of Texas Health Science Center (UTHealth). C57BL/6J, not C57BL/6N, was used as WT control because both Chi3l1^-/-^ and CD44^-/-^ mice are on the C57BL/6J background, determined by PCR (data not shown). Animal studies described have been approved by the UTHealth Institutional Animal Care and Use Committee (IACUC). For APAP treatment, mice (8-12 weeks old) were fasted overnight (5:00pm to 9:00am) before *i.p*. injected with APAP (Sigma, A7085) at a dose of 210 mg/kg for male mice and 325 mg/kg for female mice, as female mice are less susceptible to APAP-induced liver injury.[47] Male mice have been the choice in the vast majority of the studies of AILI reported in the literature.[8, 24] Therefore, we used male mice in the majority of the experiments presented. However, we observed a similar phenotype in female Chi3l1^-/-^ and CD44^-/-^ mice as in male mice (Supplementary Figure 5). In some experiments, APAP-treated mice were immediately injected intraperitoneally (*i.p*.) with either PBS (100μl) or recombinant mouse Chi3l1 (rmChi3l1, 500 ng/mouse in 100μl, Sino Biological 50929-M08H). Livers were harvested at time points indicated in the figure legends and immunofluorescence (IF) staining was performed using frozen sections to detect Mϕs and platelets using anti-F4/80 and anti-CD41 antibodies, respectively. Liver paraffin sections and sera were harvested at time points indicated in the figure legends. H&E staining and ALT measurement to examine liver injury were performed using a diagnostic assay kit (Teco Dignostics, Anaheim CA).

#### Blocking endogenous podoplanin

Mice were *i.v*. injected with Ctrl IgG (Bioxcell InvivoMab, BE0087, 100μg/mouse) or anti-podoplanin antibody (Bioxcell InvivoMab, BE0236, 100μg/mouse) in Chi3l1^-/-^ reconstituted with rmChi3l1 at 16h prior to APAP treatment.

#### Platelet depletion

WT mice were *i.v*. injected with Ctrl IgG (BD Pharmingen, 553922, 2mg/kg) or CD41 antibody (BD Pharmingen, 553847, 2mg/kg) to deplete platelets at 12h prior to APAP treatment.

#### KCs depletion

WT mice were *i.v*. injected with empty liposomes (PBS, 100μl/mouse) or clodronate-containing liposomes (CLDN, 100μl/mouse) to deplete KCs at either 9hrs or 40hrs prior to APAP treatment. Clodronate-containing liposomes were generated as previously described[24].

#### Evaluation of the effects of anti-Chi3l1 monoclonal antibodies

To examine the therapeutic potential of anti-mouse Chi3l1 mAbs, WT mice were injected (*i.p*.) with either Ctrl IgG or anti-mouse Chi3l1 antibody clones 3h after APAP administration. To examine the therapeutic potential of anti-human Chi3l1 mAbs, Chi3l1^-/-^ mice treated with APAP were immediately injected (*i.p*.) with either PBS (100μl) or recombinant human Chi3l1 (rhChi3l1, 1μg/mouse in 100μl, Sino Biological 11227-H08H). After 3h, these mice were divided into two groups injected (*i.p*.) with either Ctrl IgG or anti-human Chi3l1 mAbs.

### Bio-layer interferometry

The binding affinity between Fc-CD44 and His-Chi3l1 was measured using the Octet system 8-channel Red96 (Menlo Park). Protein A biosensors and kinetics buffer were purchased from Pall Life Sciences (Menlo Park). Fc-CD44 protein was immobilized onto Protein A biosensors and incubated with varying concentrations of recombinant His-Chi3l1 in solution (1000 nM to 1.4 nM). Binding kinetic constants were determined using 1:1 fitting model with ForteBio’s data analysis software 7.0, and the KD was calculated using the ratio Kdis/Kon (the highest 4 concentrations were used to calculate the KD.).

### Immunohistochemical (IHC) and immunofluorescent (IF)

H&E staining and IHC were performed on paraffin sections using the following antibodies: anti-human CD41 (Proteintech, 24552-2-AP, 1:200), anti-human CD68 (Thermo Fisher, MA5-13324, 1:100), anti-human Chi3l1 (Proteintech, 12036-1-AP, 1:100), and anti-mouse F4/80 (Bio Rad, MCA497R, 1:200). IF staining was performed on frozen sections using the following antibodies: anti-mouse CD41 (BD Bioscience, Clone MWReg 30), mouse F4/80 (Biolegend, 123122, 1:100), anti-CD44 (abcam, clone KM81, ab112178, 1:200), anti-Chi3l1 (Proteintech, 12036-1-AP, 1:100), anti-Podoplanin (Novus, biological, NB600-1015, 1:100), and anti-Clec-2 (Biorbyt, orb312182, 1:100). Alexa 488-conjugated donkey anti-rat immunoglobulin (Invitrogen, A-21208, 1:1000) was used as a secondary antibody for CD41 and CD44 detection. Alexa 488-conjugated goat anti-rabbit immunoglobulin (Invitrogen, A-11034, 1:1000) was used as a secondary antibody for Clec-2 detection. Alexa 594-conjugated goat anti-rabbit immunoglobulin (Invitrogen, A-11012, 1:1000) was used as a secondary antibody for Chi3l1 detection. Alexa 594-conjugated goat anti-hamster immunoglobulin (Invitrogen, A-21113, 1:1000) was used as a secondary antibody for Podoplanin detection. Nuclei were detected by Hoechst (Invitrogen, H3570, 1:10000).

### Intravital confocal microscopy

Mice were prepared for intravital microscopy as previously described.[48] Briefly, mice were anesthetized using pentobarbital and underwent tracheostomy (to facilitate breathing) and internal jugular catheterization (for antibody administration) followed by liver exteriorization as described by Marques et al,[49] with modifications. Mice were placed supine on a custom-made stage with the liver overlying a glass coverslip wetted with warmed saline and surrounded with wet saline-soaked gauze. Mice were kept euthermic at 37°C using radiant warmers and monitored with a rectal thermometer. Anesthesia was maintained using an isoflurane delivery device (RoVent with SomnoSuite; Kent Scientific) with 1-3% isoflurane delivered. Mice were intravenously injected with an antibody mixture in sterile 0.9% sodium chloride containing TRITC/bovine serum albumin (Sigma; to label the vasculature; 500 μg/mouse), BV421-anti-F4/80 antibody (to label Kupffer; 0.75 μg/mouse), and DyLight 649/anti-GPIbβ antibody (emfret analytics; to label platelets; 3 μg/mouse) for visualization. Mice were imaged on an Olympus FV3000RS laser scanning confocal inverted microscope system at 30 fps using a 60X/NA1.30 silicone oil objective with 1X and 3X optical zoom using the resonance scanner. This allows for simultaneous excitation and detection of up to four wavelengths. All animals were euthanized under a surgical plane of anesthesia at the end of the experiments.

### Image analysis of intravital microscopy experiments

The images were then analyzed by a blinded investigator to assess platelet area. Eleven to fifteen 1-minute fields of view (1X optical zoom) were analyzed per mouse using FIJI/ImageJ software. Background noise was removed using a Guassian filter (1 pixel) for all channels prior to analysis. Vascular area was measured in each field using the region of interest (ROI) selection brush in the TRITC (albumin) channel. The platelet area within the vascular ROI was then determined using threshold of the DyLight 649 (platelet) channel.

### Generation of Chi3l1 mAbs

Rabbit monoclonal antibodies were generated using previously reported methods.[50] Briefly, two New Zealand white rabbits were immunized subcutaneously with 0.5 mg recombinantly expressed human Chi3l1 protein (Sino Biological, Cat: 11227-H08H). After the initial immunization, animals were given boosters three times in a three-week interval. Serum titers were evaluated by indirect enzyme-linked immunosorbent assay (ELISA) and rabbit peripheral blood mononuclear cells (PBMC) were isolated after the final immunization. A large panel of single memory B cells were enriched from the PBMC and cultured for two weeks, and the supernatants were analyzed by ELISA. To isolate mouse Chi3l1 antibody, the rabbits were boosted twice more with mouse Chi3l1 before the memory B cell culture. The variable region genes of the antibodies from these positive single B cells were recovered by reverse transcription PCR (RT-PCR) and cloned into the mammalian cell expression vector as described previously.[50] Both the heavy and light chain constructs were co-transfected into Expi293 cell lines using transfection reagent PEI (Sigma). After 7 days of expression, supernatants were harvested and antibodies were purified by affinity chromatography using protein A resin as reported before.[50]

### Statistics

Data were presented as mean ± SEM unless otherwise stated. Statistical analyses were carried out using GraphPad Prism (GraphPad Software). Comparisons between two groups were carried out using unpaired Student t test. Comparisons among multiple groups (n>=3) were carried out using one-way ANOVA. P values are as labeled and less than 0.05 was considered significant. Platelets counts/mm^2^ was analyzed by Image J software.

### Study approval

Serum samples from patients diagnosed with APAP-induced liver failure on day 1 of admission were obtained from the biobank of the Acute Liver Failure Study Group (ALFSG) at UT Southwestern Medical Center, Dallas, TX, USA. The study was designed and carried out in accordance with the principles of ALFSG and approved by the Ethics Committee of ALFSG (HSC-MC-19-0084). Formalin-fixed, paraffin-embedded human liver biopsies from patients diagnosed with APAP-induced liver failure were obtained from the National Institutes of Health-funded Liver Tissue Cell Distribution System at the University of Minnesota, which was funded by NIH contract # HHSN276201200017C.

See Supplementary Material for details for other methods.

## Author contributions

ZS designed and conducted the experiments, analyzed/interpreted the data, and wrote the manuscript; LL generated anti-human/mouse Chi3l1 antibodies; CLA conducted experiments and analyzed data; XG performed the interferometry assay; FWL conducted the intravital microscopy experiments; DCF and BG provided slides of healthy individuals and patients with AILI; LW provided patients’ serum samples; BG, YW, JJ, NFM and ZQA revised manuscript and provided suggestions; CGL and JAE provided Chi3l1^-/-^ mice; ZQA and NYZ supervised the generation of anti-human/mouse Chi3l1 antibodies; CJ conceived and supervised the project and wrote the manuscript.

## Acknowledgements

We appreciate the time and effort from Dr. Yanyu Wang (Department of Anesthesiology, UTHealth), who diligently double-checked the raw data for each figure. ZS received funding from NSFC (32071129). FWL received funding from NIH (GM123261). ZA received funding from CPRIT (RP150551 and RP190561) and the Welch Foundation (AU-0042-20030616). C.J. received funding from NIH (DK122708, DK109574, DK121330, and DK122796) and support from a University of Texas System Translational STARs award. Portions of this work was supported with resources and the use of facilities of the Michael E. DeBakey VA Medical Center and funding from Department of Veterans Affairs I01 BX002551 (Equipment, Personnel, Supplies). The contents do not represent the views of the U.S. Department of Veterans Affairs or the United States Government.

## Supplementary Materials and Methods

### Blocking endogenous CD44

Mice were *i.p*. injected with Ctrl IgG (BD Pharmingen, 559478, 50μg/mouse) or anti-CD44 antibody (BD Pharmingen, 553131, 50μg/mouse) in Chi3l1^-/-^ reconstituted with rmChi3l1 at 30 min prior to APAP treatment.

### Preparation of liver cells and *in vitro* cell culture

Hepatic nonparenchymal cells (NPCs) and hepatocytes were isolated as previously described[20]. In brief, mice were anesthetized and liver tissues were perfused with EGTA solution, followed by a 0.04% collagenase digestion buffer. Liver hepatocytes and NPCs were isolated by gradient centrifugation using 35% percoll (Sigma). To further purify LSEC and Mϕs, LSEC and Mϕs fractions were stained with phycoerythrin (PE)-conjugated anti-CD146(for LSEC, Invitrogen, 12-1469-42), and anti-F4/80(for Mϕs, Invitrogen, 12-4801-82) antibodies and positively selected using EasySep™ Mouse PE Positive Selection Kit (Stemcell technologies) following manufacturers’ instructions. Each subset will yield a purity around 90%.

#### Co-culture of Mϕs and platelets

Isolated Mϕs were cultured in DMEM with 10% fetal bovine serum and pre-treated with Podoplanin antibody (Bioxcell InvivoMab, BE0236, 2μg/ml) for 30mins and then co-culture with washed platelets for 30mins. Unbound platelets were washed out and Podoplanin and Clec-2 on Mϕs were stained.

### Isolation of platelets

Mouse whole blood was collected with anti-coagulant ACD solution from Inferior vena cava. Platelets were further isolated by additional washes with Tyrode’s buffer. Isolated washed platelets were subjected to functional assay after incubation with PGI_2_ (Sigma, P6188) for 30mins.

### Flow cytometry

Isolated liver NPCs were incubated with1μl of anti-mouse FcγRII/III (Becton Dickinson, Franklin Lakes, NJ, USA) to minimize non-specific antibody binding. The cells were then stained with anti-mouse CD45-V655 (eBioscience, 15520837), F4/80-APC/Cy7 (Biolegend, 123118), Ly6C-APC (BD Pharmingen, 560595), Ly6G-V450 (BD Pharmingen, 560603), CD146-PerCP-Cy5.5(BD Pharmingen, 562134), CD44-PE (BD Pharmingen, 553134), anti-His-FITC (abcam, ab1206). In some experiments, cells were incubated with 2μg rmChi3l1 for 2h before antibody staining. The cells were analyzed on a CytoFLEX LX Flow Cytometer (Beckman coulter, IN, USA) using FlowJo software (Tree Star, Ashland, OR, USA). For flow cytometric analysis, CD45^+^ cells were gated to exclude endothelial cells, hepatic stellate cells, and residue hepatocytes. Within CD45^+^ cells, CD44^+^ cells that bind to Chi3l1 were back gated to determine the cells types.

### Extraction of liver proteins, immunoprecipitation, and mass spectrometry

Snap frozen liver tissues were pulverized to extract liver proteins in STE buffer. Protein concentration was measured by BCA kit (Thermo Scientific, 23225) following the manufacturer’s instructions.

#### Immunoprecipitation of NPCs lysates

Proteins were extracted from NPCs lysates and incubated with 5μg rmChi3l1, followed by immunoprecipitation with 2ug Rabbit IgG (negative control, Peprotech, 500-p00) or 2μg anti-his tag antibody (Abnova, MAB12807). Dynabeads Protein G (Invitrogen, 1003D) were used to pull down antibodies-binding proteins. Immunoprecipitated proteins were subject to mass spectrometry analyses by the Proteomics Core Facility at UTHealth.

#### Immunoprecipitation of liver homogenates

CD44^-/-^ and WT mice were treated with APAP for 2h. 10mg liver proteins were extracted from treated mice and incubated with 5μg rmChi3l1, followed by immunoprecipitation with 2μg anti-CD44 antibody (BD Pharmingen, 553131). Dynabeads Protein G (Invitrogen, 1003D) were used to pull down antibody-binding proteins. Input and immunoprecipitated proteins were subject to western blot analyses.

#### In vitro immunoprecipitation assays

2μg rhChi3l1(Sino Biological, His Tag, 11227-H08H) or 2μg GST protein (His Tag) as control were incubated with 2μg human CD44 (Sino Biological, Fc Tag, 12211-H02H) and immunoprecipitated with 2μg anti-His antibody (Abnova, MAB12807). Input and immunoprecipitated proteins were subject to western blot analyses.

### Western blotting

Samples were prepared with loading buffer and boiled before loading onto SDS-PAGE gels. Nitrocellulose membranes (Bio-Rad) were used to transfer proteins. Primary antibodies used to detect specific proteins: anti-Chi3l1 (Proteintech, 12036-1-AP, 1:1000), anti-CD44 (abcam, ab25340, 1:500), anti-β-actin(Cell Signaling, 4970, 1:1000), anti-His (Abnova, MAB12807, 1:1000), anti-cyp2e1 (LifeSpan BioSciences, LS-C6332, 1:500), anti-APAP adducts[24] (provided by Dr. Lance R. Pohl, NIH, 1:500). Secondary antibodies include goat anti-Rabbit IgG (Jackson ImmunoResearch, 111-035-144, 1:1000), goat anti-Rat (Jackson ImmunoResearch, 112-035-003, 1:1000).

### Quantitative Real-Time Reverse Transcriptase Polymerase Chain Reaction (qRT-PCR)

Total RNA was isolated from 1×10^6^ cells using RNeasy Mini Kit (Qiagen, Valencia, CA). After the removal of genomic DNA, RNA was reversely transcribed into cDNA using Moloney murine leukemia virus RT (Invitrogen, Carlsbad, CA) with oligo (dT) primers (Invitrogen). Quantitative PCR was performed using SYBR green master mix (Applied Biosystem) in triplicates on a Real-Time PCR 7500 SDS system and software following manufacturer’s instruction (Life Technologies, Grand Island, NY, USA). RNA content was normalized based on amplification of 18S ribosomal RNA (rRNA) (18S). Change folds = normalized data of experimental sample/normalized data of control. The specific primer pairs used for PCR are listed in Table 1 below:

**Table 1.**
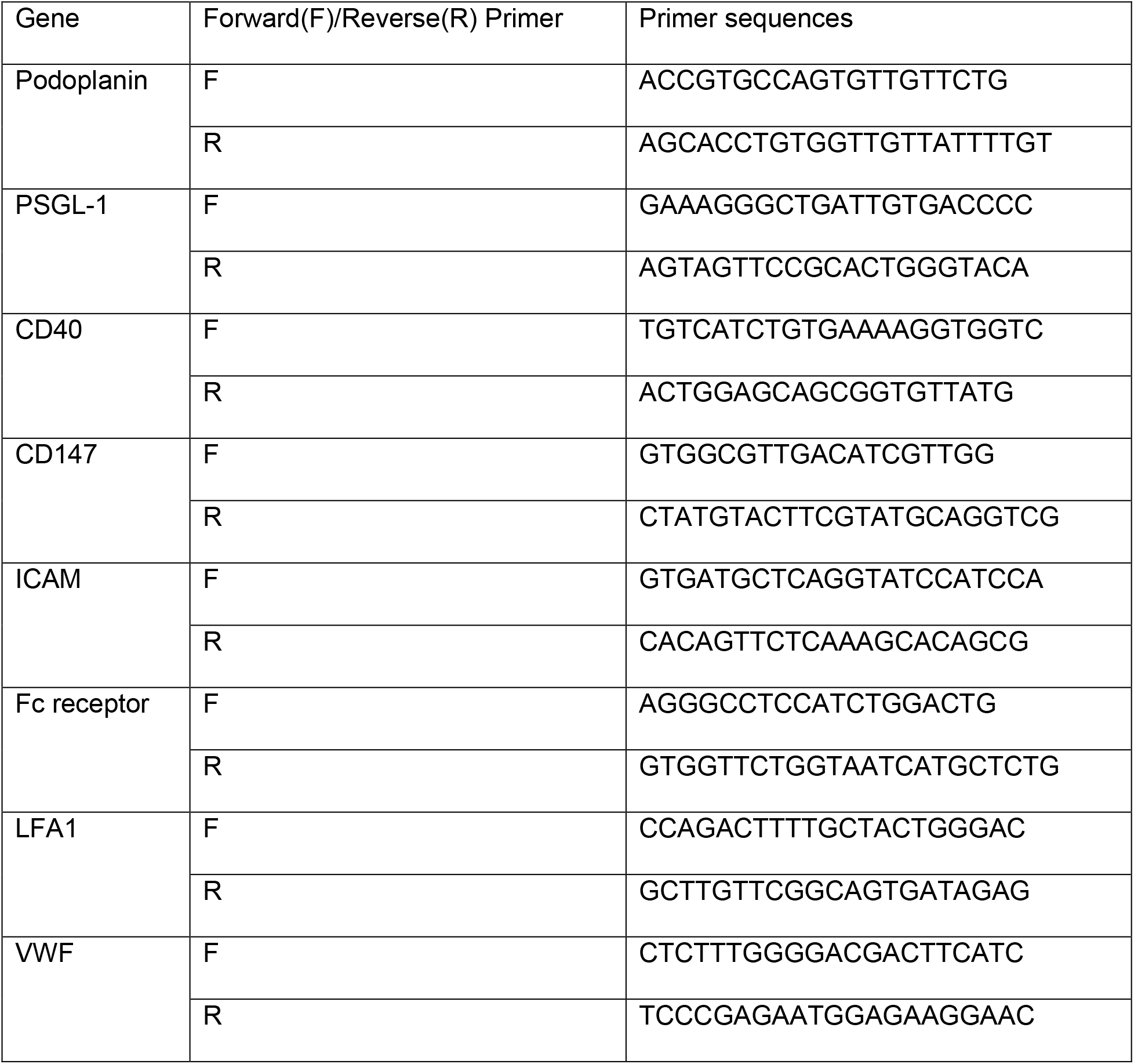
Real-Time PCR Primers used

## Supplementary Figures

**Supplementary Figure 1.**
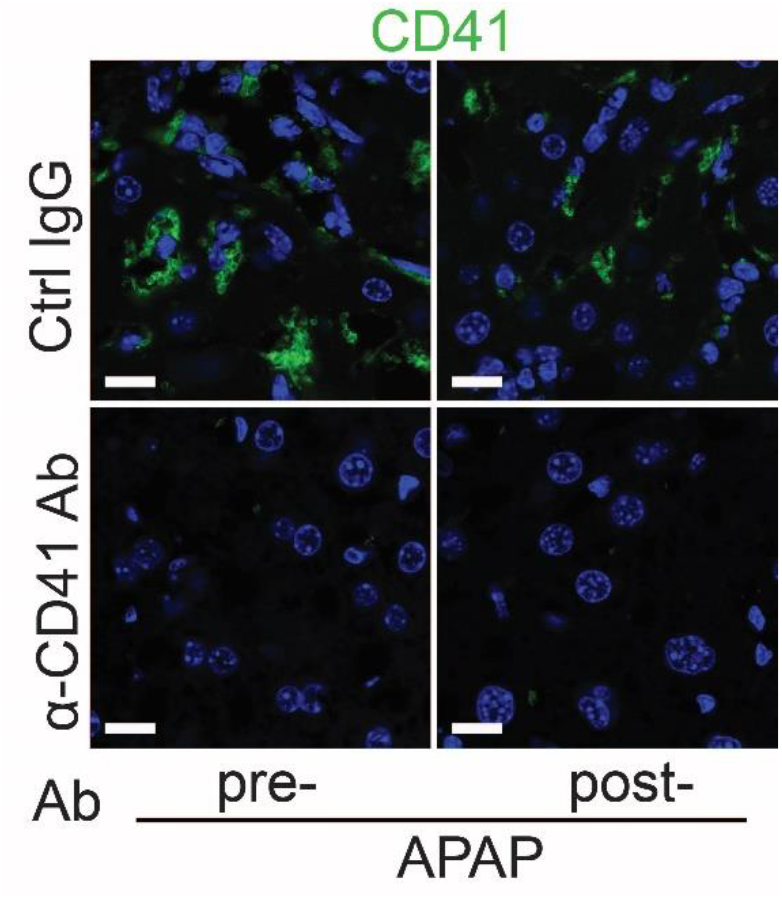
Depletion of platelets by anti-CD41 antibody reduces hepatic platelets recruitment. Male C57B/6 mice were treated with control IgG (Ctrl IgG) or an anti-CD41 antibody (α-CD41 Ab) either 3hrs before (pre-) or 3hrs after (post) APAP administration. IF staining was performed to identify intrahepatic platelets (CD41^+^) (n=5 mice/group). Scale bar, 25μm.

**Supplementary Table 1.**
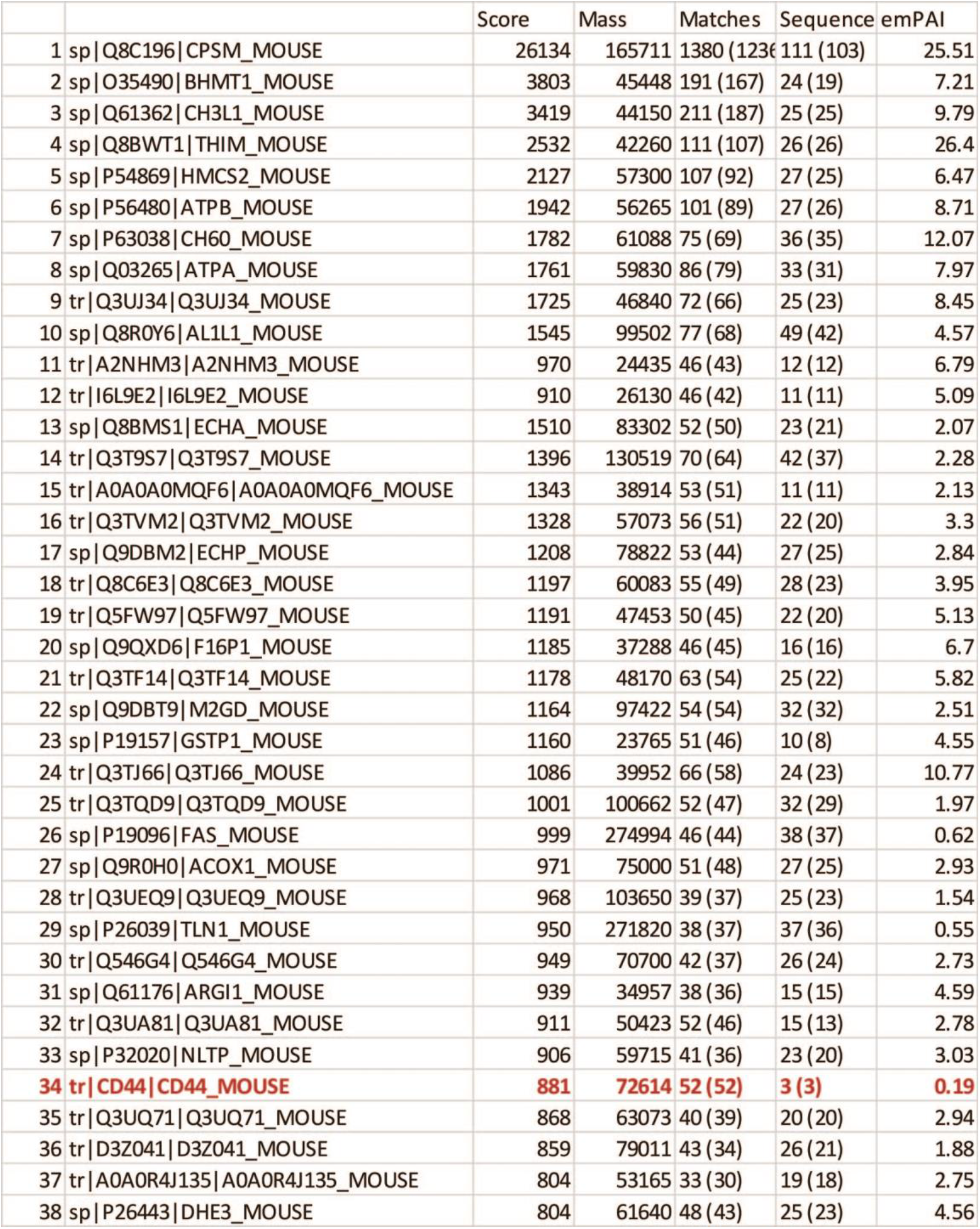
Representative list of potential Chi3l1-interacting proteins detected by mass spectrometry. Non-parenchymal cells were isolated from C57B/6 mice treated with APAP for 3hrs and the cell lysate was incubated with rmChi3l1 overnight. Proteins potentially bound to rmChi3l1 were immune-precipitated with an anti-His antibody and subjected to mass spectrometry analyses.

**Supplementary Figure 2.**
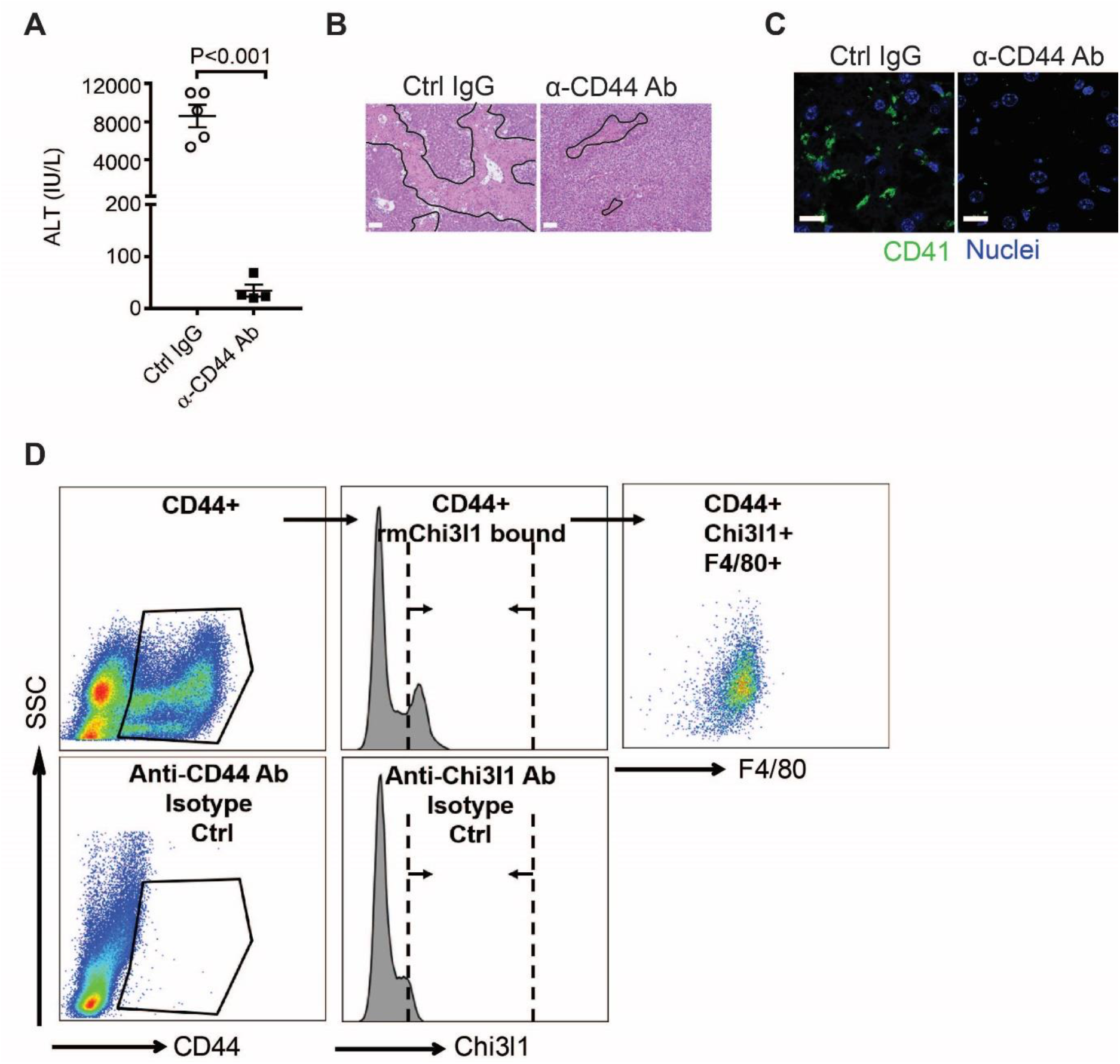
Chi3l1 promotes hepatic platelet recruitment and AILI through CD44 expressing on Mϕs. (**A-C**) Chi3l1^-/-^ mice reconstituted with rmChi3l1 were treated with either Ctrl IgG or α-CD44 Ab 30 min prior to APAP challenge. (**A**) Serum levels of ALT and (**B**) liver histology with necrotic areas outlined were evaluated 24hrs after APAP treatment (n=4-5 mice/group). Scale bar, 250μm. (**C**) IF staining was performed to detect intrahepatic platelets (CD41^+^) 3hrs after APAP treatment (n=3 mice/group). Scale bar, 25μm. Two-tailed, unpaired student t-test was performed in **A**. (**D**) Flow cytometry analysis was performed to identify Chi3l1-binding cells among liver non-parenchymal cells (NPCs) isolated from WT mice treated with APAP for 2hrs. CD44^+^ cells were gated from single live cells. CD44^+^ cells that bind to rmChi3l1 were further gated. The Chi3l1^+^CD44^+^ cells were then identified by markers for various cell types, including CD45^+^ CD146^-^F4/80^+^(Mϕs), CD45^-^CD146^+^(LSECs) and Ly6G^+^(neutrophils).

**Supplementary Figure 3.**
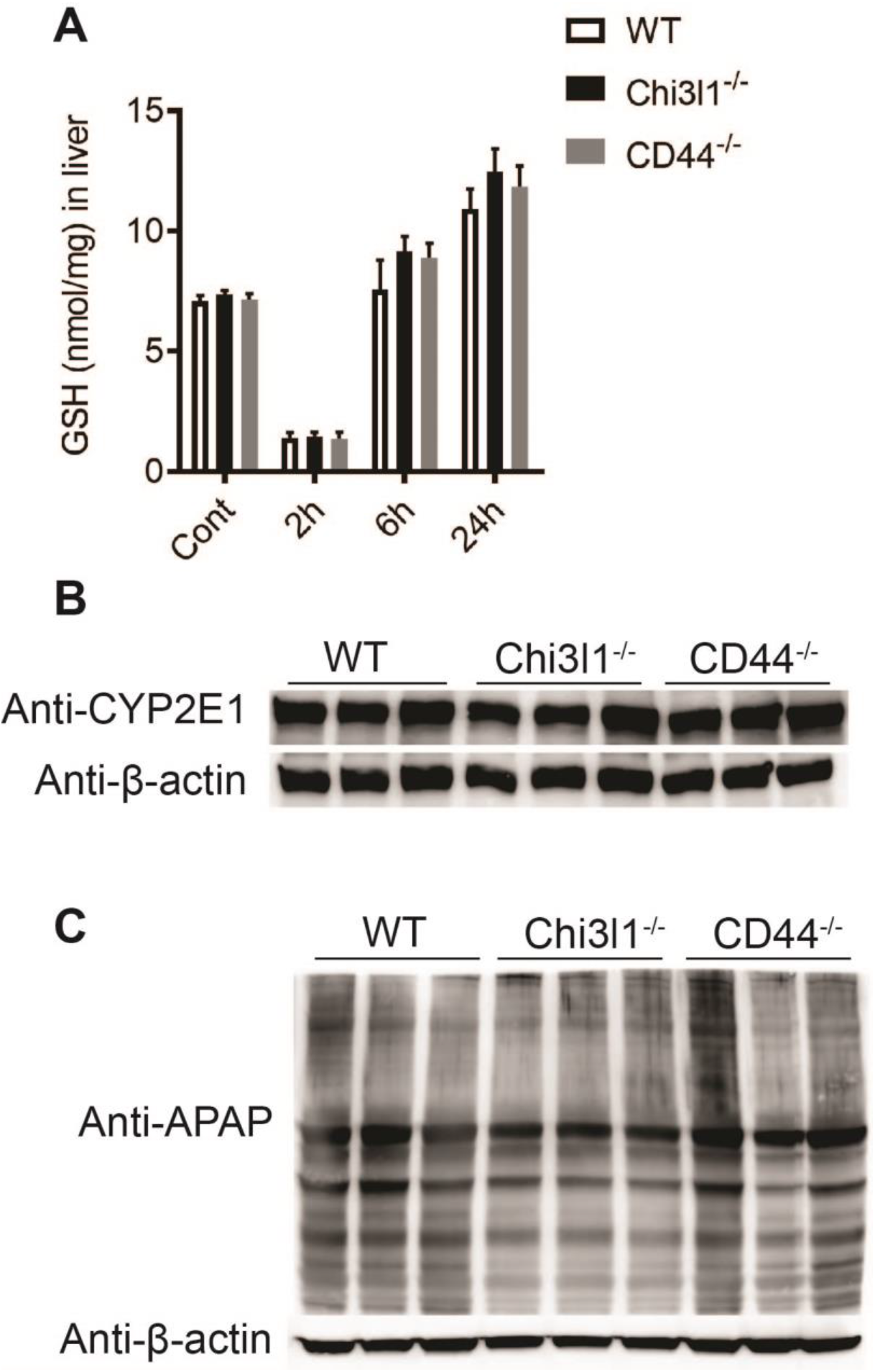
Deletion of Chi3l1- nor CD44 affects APAP bio-activation. Male C57B/6 mice were treated with APAP (n= 3 mice/group). (**A**) GSH levels in the liver were measured at indicated time points by HPLC. (**B**) Hepatic protein levels of CYP2E1 were measured by Western blotting after mice were fasted overnight without APAP treatment. (**C**) NAPQI-protein adducts in liver were measured by Western blotting 2hrs after APAP treatment.

**Supplementary Figure 4.**
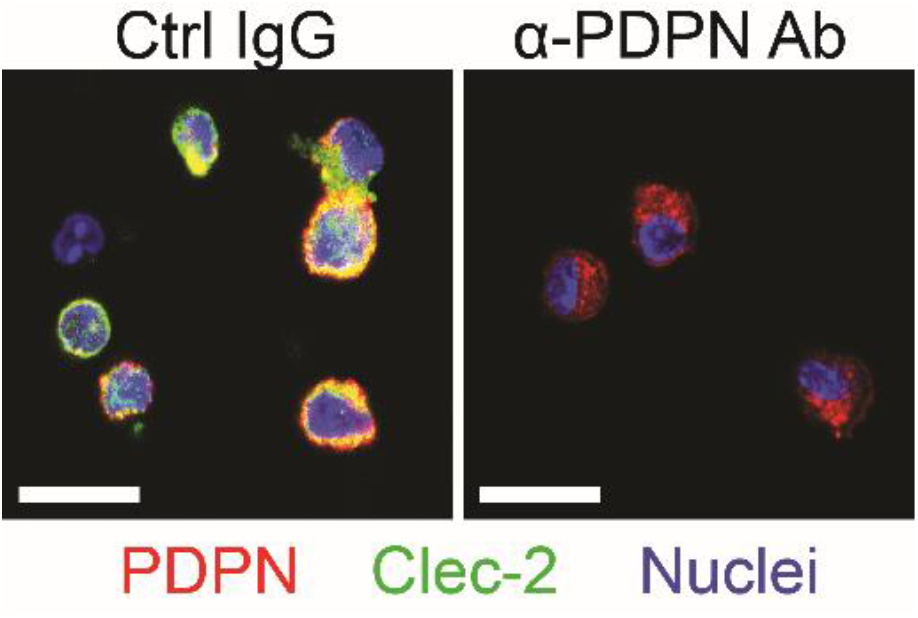
Podolanin expressing on Mϕs mediates interactions with platelets. Mϕs were isolated from WT mice treated with APAP for 3hrs. The cells were treated *in vitro* with either control IgG (Ctrl IgG) or an anti-podoplanin antibody (α-PDPN Ab) before incubation with platelets. IF staining was performed to detect PDPN on Mϕs and Clec-2 on platelets. Scale bar, 25μm.

**Supplementary Figure 5.**
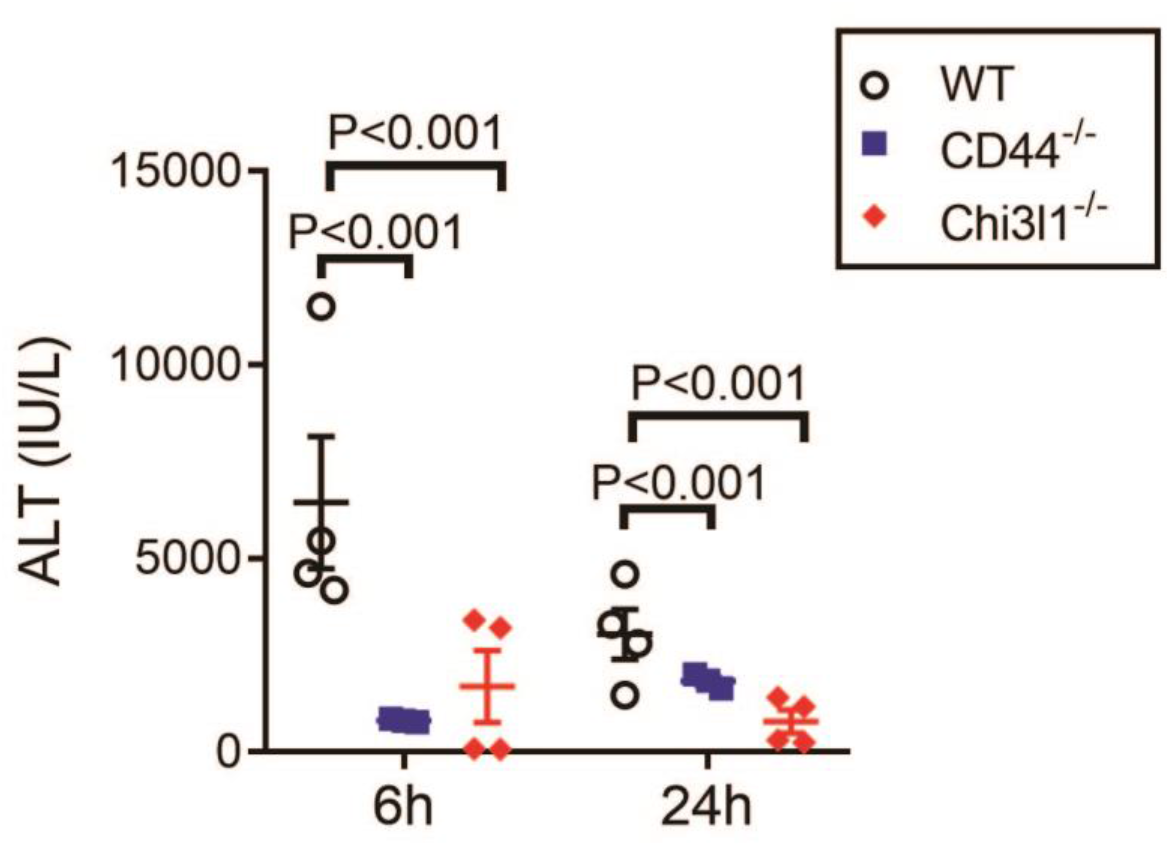
Female Chi3l1^-/-^ and CD44^-/-^ mice develop reduced liver injury compared to female WT mice. Female WT, Chi3l1^-/-^ and CD44^-/-^ mice were treated with APAP. Serum ALT levels were measured at 6hrs and 24hrs after APAP treatment (n=6-8 mice/group). One-way ANOVA was performed.

